# Cryo-EM structure of the diapause chaperone artemin

**DOI:** 10.1101/2022.04.06.487001

**Authors:** Amar D. Parvate, Samantha M. Powell, Jory T. Brookreson, Trevor H. Moser, Irina V. Novikova, Mowei Zhou, James E. Evans

## Abstract

The protein artemin constitutes over 10% of all protein in *Artemia* cysts during diapause and acts as both an RNA and protein chaperone. However, its mechanistic details remain elusive since no high-resolution structure of artemin exists. Here we report the full-length structure of artemin at 2.04 Å resolution. The cryo-EM map contains density for an intramolecular disulfide bond between Cys22-Cys61 and resolves the entire C-terminus extending into the core of the assembled protein cage. We also provide data supporting the role of C-terminal helix F towards stabilizing the dimer form that is believed to be important for its chaperoning activity. We were able to destabilize this effect by placing a tag at the C-terminus to fully pack the internal cavity and cause limited steric hindrance.

## 1 Introduction

Species of the brine shrimp *Artemia* are found across North, Central and South America and inhabit some of the most challenging environments^1^. The key to surviving such harsh conditions has been tracked to the brine shrimp’s ability as a cyst to enter a state of metabolic hypoactivity called diapause. In this state, the cyst can survive desiccation, high and low temperatures, radiation and years of anoxia ^2^. A complement of stress tolerance proteins have been reported in *Artemia* during diapause including p26, artemin and hsc70 ^1^. Of this group, artemin is particularly interesting due to evidence that it acts as both a protein and RNA chaperone ^3^. Excluding the yolk, artemin can constitute 10-15% of the total protein content of cysts in diapause ^4^. Additionally, *in vitro* studies have shown artemin to be highly thermostable and to demonstrate chaperone-like activity under prime stressors such as exposure to heat, H_2_O_2_, or both, and also exposure to cold ^4-6^.

While artemin is a ferritin homolog, its differences rather than similarities to ferritins shed more light on its role as a chaperone. Artemin monomers are 229 amino-acid residues long with a molecular mass of 26 kDa and the 24mer has a mass of ∼624 kDa. The artemin monomer is 45–50 residues longer than most ferritins, even though they form oligomers of similar dimensions and symmetry ^7^. Unlike ferritins whose job is to sequester iron, artemin is unable to bind iron due to naturally modified regions of the ferroxidase center, iron nucleation center and 3-fold channel. Additionally, artemin is a thiol rich molecule with 9 free thiols and one thiol involved in a disulfide bond ^8^. Importantly, several biochemical studies point to the chaperone activity of artemin being regulated by a redox switch courtesy of the thiols ^9^ as well as its C-terminus which diverges considerably from ferritins ^10^.

All prior structural hypotheses for artemin function were based upon computationally derived homology models using apoferritin as a template ^11^. The homology models indicated that the core of artemin has a similar fold to apoferritin, including the 5 core ferritin helices (A-E) and the hydrophobic loop L. However, the first twenty N-terminal residues of artemin were suggested to exist as flexible loops directed outwards and solvent exposed, while the C-terminal residues were predicted to curve inwards into the cavity of the artemin ^12^. Other than *in silico* data suggesting that the C-terminus completely fills the central cavity of artemin, there was no consensus in prior literature on the fold or secondary structure of the C-terminus despite this region having significant roles in chaperone activity. Additionally, none of the prior reported homology models are currently publicly available as they were not posted to sustained repositories and this makes continued studies difficult.

Based on homology models and biochemical data, a mechanism of action for the chaperoning activity of artemin has been suggested to rely on the activation through a cysteine redox switch in response to environmental stressors. This leads to the 24mer breaking down into oligomers of which dimers are believed to be most abundant and the functional chaperone ^13^. The stable dimer putatively interacts with the target protein through the C-terminal helices to stabilize the target protein and prevent either denaturing or unfolding or both. Chaperone activity has been observed to stay at peak levels under multiple conditions such as between 25-50 °C, in presence of 40-100 mM hydrogen peroxide, and following exposure to cold or hypersaline environments ^5,14^. Several factors have been proposed to play an essential role in artemin chaperoning activity including the number of free and solvent exposed thiols, existence of exposed hydrophobic surfaces and also the local environment of Trp, Tyr and His residues ^8,9^. However, the absence of a high-resolution structure of artemin has led to competing theories for artemin’s mechanism of action based on prior homology models, and the ultimate structural details of the protein elusive.

Here we used an integrative approach combining cell-free expression, cryo-electron microscopy and native mass spectrometry to determine the atomic structure of artemin. We provide a structure of full-length artemin at 2.04 Å using single particle cryo-EM coupled with cell-free expression. Native mass spectrometry (MS) was used to confirm the molecular weight of all species and probe the stability of artemin dimerization since the dimer form is believed to be the functional subunit while chaperoning.

## 2 Materials and Methods

### 2.1 Protein expression and purification

DNA plasmids for artemin were prepared by Genscript using their custom gene synthesis and cloning services. Obtained DNA templates (pEU_artemin_6His and pEU_3XF_artemin) were used in the cell-free gene expression and protein purification by Protemist DTII, an automated protein synthesizer from CellFree Sciences, using well-established in-house protocols ^15^ and manufacturer’s guidelines. **Supplementary Table 1** shows the amino acid sequences for all clones. For 3XFLAG-based purification on the Protemist DTII, 800 µl of ANTI-FLAG M2 Affinity gel (Sigma, A2220) was used per 6-ml translation reaction. In addition, in all reactions, SUB-AMIX buffer was supplemented with protease inhibitor cocktail (Sigma Aldrich, #539137) with the buffer to cocktail ratio of 100:1 (v/v). For the expression of fluorophore-labeled proteins, the translation mixture was supplemented with FluoroTect GreenLys reagent (Promega). Purified samples were washed with TBS (50mM Tris and 150mM NaCl, pH 7.5) buffer and concentrated in a pre-chilled centrifuge at 15,000×g to a final volume of 500 µL using a 0.5 mL 10kDA MWCO spin column Concentrated proteins were further loaded onto an AKTA Pure system stored at 4°C using either a Superose 6 Increase 10/300 or Superdex 200 Increase 10/300 column. Aliquots within a peak on the AKTA SEC trace were combined and concentrated using 10 kDa MWCO Amicon spin columns to a final volume of 100 µL. Protein purity was verified by SDS and Native PAGE.

### 2.2 Cryo-EM sample preparation and single particle data collection

Three µL of artemin solution at 0.2-1.5 mg/ml were loaded on to glow discharged Quantifoil grids (200 mesh R2/1 or 300 mesh R1.2./1.3). Grids were blotted for 1.5-3.5 s and plunge frozen in liquid ethane on a Leica EM GP2. Grids were stored in liquid nitrogen until further use. For screening and data collection, grids were loaded on a 300 keV Titan Krios G3i (Thermo Fisher) and all datasets were collected using the standard EPU software along with K3 direct electron detector and a Bioquantum energy filter (Gatan Inc) with 20 eV slit. Movies were collected at 130,000× magnification in super resolution mode resulting in a pixel size of 0.3398 Å respectively. Movies were collected at a total dose ranging from 41.7 to 58.9 e^-^/Å^2^, with 0.5 to 1.8 s exposures, and a defocus range between - 0.3 to -1.3 µm.

### 2.3 Image processing

All the movies were processed using cryoSPARC Live and cryoSPARC ^16^. Motion correction and CTF estimation were performed using default parameters and initial particle extraction used the built-in *blob picker* with a box size of 400 or 800 pixels ^17^. Details about particle numbers at each step are listed in **Supplementary Table 2 and Supplementary Figure 1**. Initial subsets of particles were subjected to reference free 2D classification before discreet and diverse classes were chosen to re-extract particles using template picking. Multiple rounds of classification were performed to exclude junk and non-homogenous classes. Ab-initio models were generated using a subset of these particles and C1 symmetry. The entire particle set was refined in 3D against ab-initio models without symmetry. Octahedral symmetry was imposed in subsequent rounds of refinement. Per particle local CTF refinement was performed before the final round of homogenous refinement. Resolution of the final map was estimated using the gold standard at 0.143 FSC. Maps were visualized using UCSF Chimera ^18^ and have been deposited in the EMDB (emdataresource.org, Flag-artemin = EMD-24706, artemin-His = EMD-24707).

**Figure 1.**
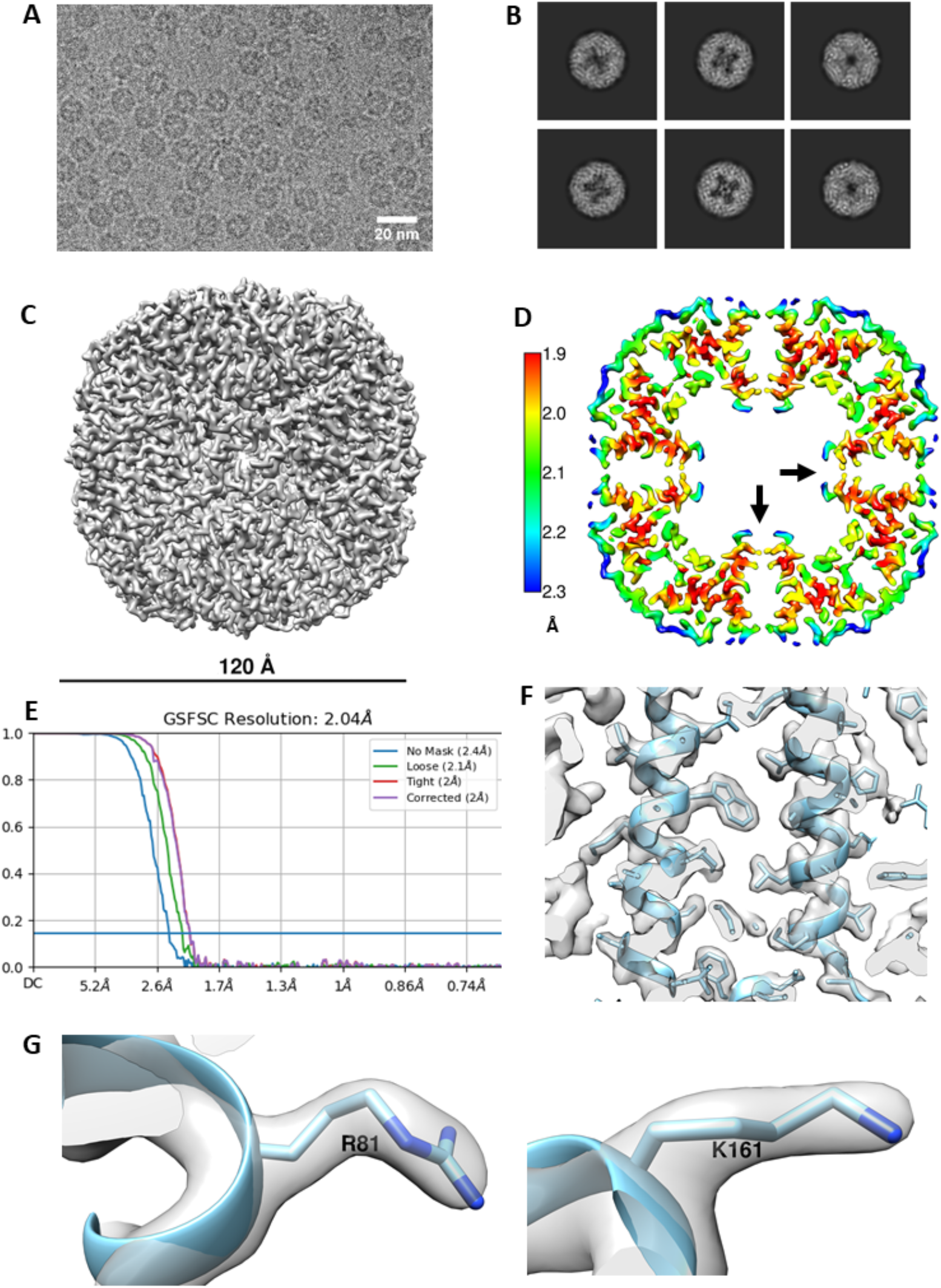
Data processing results for Flag-artemin. A) Representative micrograph of artemin showing overall dimensions similar to apoferritin but the central cavity is partially filled with density B) 2D classes showing that while the C-terminus of the monomer does point inwards from the shell, it does not fully fill the central cavity. C) Cryo-EM map of Flag-artemin with ∼120 Å diameter. D) Thin virtual slice through a resolution heat map showing the C-term alpha helices pointing inwards (black arrow) into the cavity. Scalebar indicates resolution in Å. E) Resolution estimated by gold standard at 2.14 Å at 0.143 FSC. F) Quality of the map as inspected by fitting of alpha helices and side chains of residues. G) Fitting of Arg81 and Lys161 side chains in the density.

### 2.4 Modelling

Initial homology model for artemin was generated using ^19^ HHPRED and MODELLER ^20^ based on the top 15 aligned sequences to known ferritin structures(**Supplementary Figure 2**). Models were also generated using AlphaFold2 and RosettaFold. To improve the clarity of the density map of Flag-artemin, the *Autosharpen map* tool in Phenix was used. All models were initially docked into the raw artemin map using *Dock in Map* ^21^. This was followed by an initial round of refinement with Phenix *Real-space Refinement* on the initial model from MODELLER as this had the best initial score following docking. The initial docked model was missing N-terminal residues 1-25 and C-terminal residues 202-228. Using the sharpened map, iterations of model building in COOT ^22^ and refinement in Phenix, the entirety of the C-terminus, and residues 22-25 were built into the model. Model validation of the monomer and dimer was performed using Molprobity ^23^. Using the symmetry file generated by *Map Symmetry* in Phenix, the full artemin 24-mer was modeled into the map. The final model of Flag-artemin was deposited to the PDB (https://www.rcsb.org/) (PDB: 7RVB).

**Figure 2.**
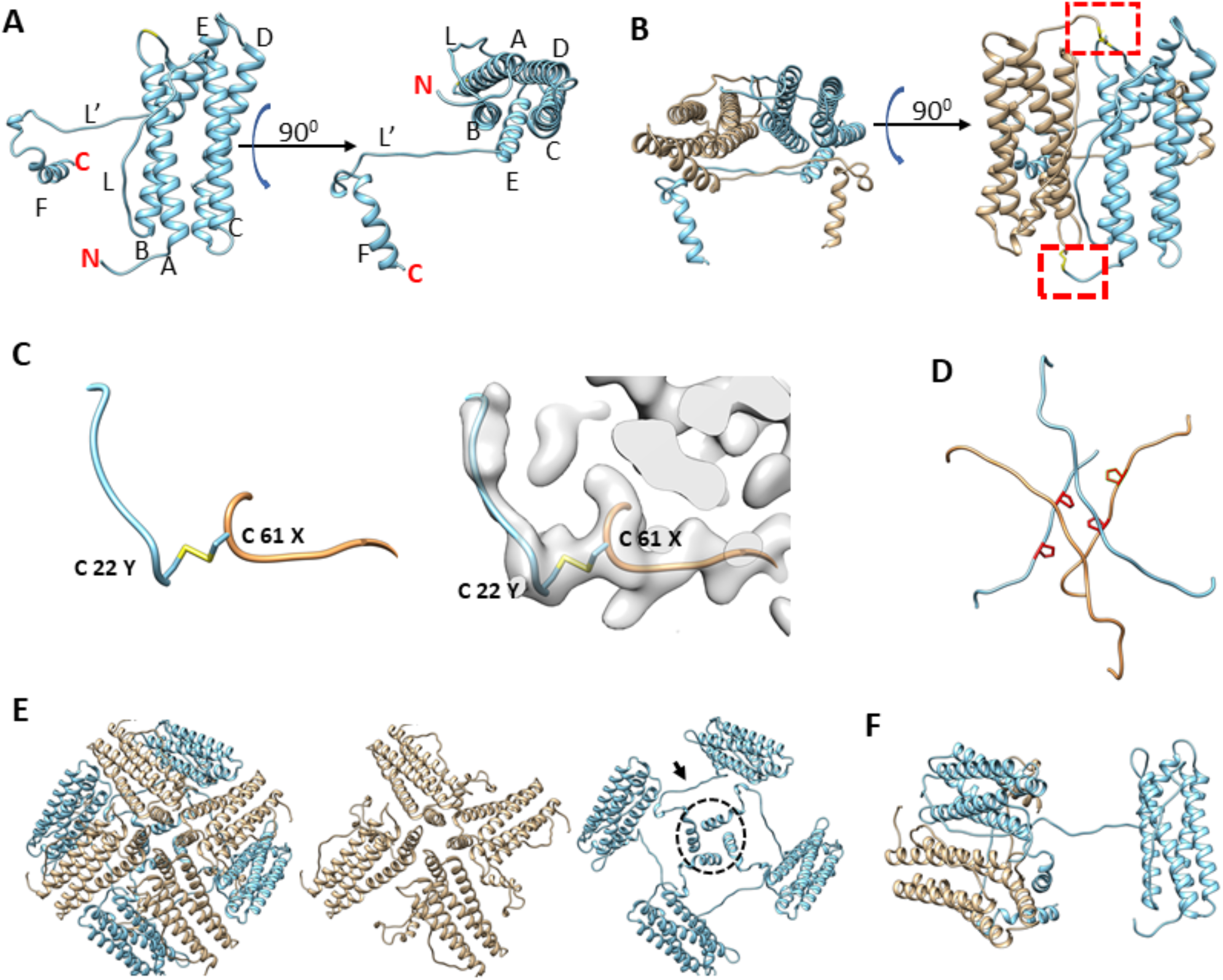
Structural organization of artemin. A) Artemin monomer with helices A-F and loops L and L’ annotated. The extra length of helix E for artemin compared to apoferritin helps position loop L’ to run along the inside of the core shell of artemin before helix F turns inward into the artemin cavity. B) Artemin dimer with antiparallel monomers colored separately (tan vs sky blue). Dashed border indicates area of the Cys22-Cys61 disulfide bond C) A zoomed in view of the region in the dashed boundary in B) shows the Cys22-Cys61 disulfide bond, with and without the map density. D) The L and L’ loops from respective monomers forming the hashtag arrangement. E) Four artemin dimers around a 4-fold axis. The conventional 4-fold axis has loops from monomers containing Cys172 (tan) similar to apoferritin arrangement. In addition, the complementary monomers (sky blue) in apoferritin have a second arrangement where loop L’ and helix F contact extend towards the neighboring dimer. F) Helix F interacts with both monomers from the neighboring dimer.

### 2.5 Native Mass Spectrometry

Protein samples were dialyzed overnight in 200 mM ammonium acetate using 96-well Microdialysis units (10k MWCO, Pierce). If further salt removal was needed, additional buffer exchange was performed using Zeba Spin Desalting Columns (7k MWCO, 75 µl, Thermo Fisher). Final concentrations used for native mass spectrometry were 1-2 µM. All native MS data was acquired on a Waters Synapt G2s-i ion mobility time-of-flight mass spectrometer. Nanoelectrospray voltage (0.6-0.8 kV) was applied through a Pt wire inserted into hand-pulled borosilicate glass capillaries (Sutter Instrument) which contained the protein solution. To filter the artemin 24-mer from low *m/z* species prior to collision induced dissociation (CID), a manual fixed quad profile of 10,000 was used. MassLynx v4.1 (Waters) was used to manually analyze spectra and mass deconvolution was performed using UniDec version 4.3.0 ^24^.

## 3 Results

### 3.1 Artemin single particle cryo-EM map revealed a unique central cavity

Ever since artemin’s first report in 1980 ^25^, a growing body of reports have elucidated the role of artemin as a molecular chaperone ^4,5,12^, but structural information about the protein had been limited to *in silico* modelling and some spectroscopic studies to date ^2^. We sought to determine the full-length structure of artemin experimentally using single particle cryo-EM. To generate the protein sample, we employed cell-free protein expression (coupled transcription and translation reactions in a test tube) and purification protocols well-established in-house (Novikova et al, 2018; Novikova et al, 2021). Using an N-terminal tagged 3XFLAG artemin construct (Flag-artemin) from *A. fransciscina* we obtained 250 µg of protein, which was sufficient quantity and purity for our needs. To obtain even more homogenous sample, artemin was further purified using size exclusion chromatography **(Supplementary Figure 1 A, B)** prior to plunge freezing on cryo-EM grids followed by single particle screening and data collection. The motion corrected cryo-EM images showed rosette-like artemin particles with a diameter of ∼120 Å **(Figure 1 A)**. However, the central cavity of artemin is not completely filled as suggested by previous modelling studies, as evident in the raw images as well as 2D class averages **(Figure 1 B)**. While no symmetry was applied for 3D ab-initio model generation and initial 3D refinement, those results clearly revealed an octahedral symmetry which matched with the expected 24mer assembly state for artemin. Therefore, octahedral symmetry was imposed in subsequent steps of 3D reconstruction and refinement and led to a final map at 2.04 Å at 0.143 FSC **(Figure 1 C-F)**.

### 3.2 Atomic model of artemin provided novel structural details

Prior sequence alignment, homology modeling and molecular dynamics studies had predicted that the structure of artemin would be similar to apoferritin ^11^ with the exception of the artemin c-terminus filling the inner cavity. After fitting an initial homology model of artemin into the cryo-EM density map the model was corrected and refined with a combination of COOT and Phenix **(Supplementary Tables 2 & 3)**. In total, all residues for artemin except the first 21 N-terminal residues were modeled and the fit confirms that the overall organization of artemin is analogous to apoferritin with residues 29 - 173 of artemin forming a similar shell structure as apoferritin (PDB: 4V1W) comprised of 5 α-helices (A-E) and one long disordered loop (L). Major differences arise due to artemin having a 28 residue long disordered N-terminus region as well as an additional helix (F) and a second long disordered loop (L’) **(Figure 2 A)**. Importantly, this experimentally determined 3D structure of artemin has density corresponding to the entire C-terminus and clearly shows that the internal cavity is not completely filled in contrast to prior in silico models. Although prior molecular dynamics simulation studies suggested that the C-terminus of artemin forms α-helices that extend inwards into the cavity of the molecule to fill the space, our cryo-EM map of Flag-artemin and the corresponding fitted atomic model clearly show that the C-terminal residues do ultimately turn inwards but they first hug the inner surface of the core artemin shell before extending only partly into the artemin cavity. Interestingly, the unique loop L’ of artemin is oriented orthogonal to loop L and the apoferritin like 4-helix bundle at the 4-fold channel. Artemin’s loop L’ contains Pro198 and Pro201 which potentially prevents this region from getting ordered into a helical conformation and helping favor the interaction with the inner surface of the shell. A third proline in the C-terminus (Pro213) provides a kink that results in the C-terminal helix F turning into the cavity of artemin (aa 216-229). **(Figure 2 A, B)**.

Previous publications noted that artemin only retains 1 of 7 conserved residues related to ferritin feridoxase activity, 1 of 6 conserved residues for the 3-fold channel and 0 of 4 conserved residues for iron nucleation ^10^. The atomic model based on the cryo-EM map clearly confirms a lack of a charged 3-fold channel. Interestingly, the residues typically associated with iron nucleation in ferritin are directly occluded in the experimentally determined artemin atomic model due to the presence of the extra loop L’. While the mutations of the four glutamate residues typically associated with iron nucleation in ferritin to Trp, His, Val and Gln in artemin would prevent nucleation simply due to the change in electrostatics, this change also facilitates the interaction with loop L’ by removing the highly negatively charged four glutamate residues in an 8-residue span. Thus, the presence of loop L’ also prevents iron nucleation. Other amino acid differences between artemin and ferritin show a general change in electrostatic surface potential even though the coulombic surface map looks very similar (**Supplementary Figure 3**).

**Figure 3.**
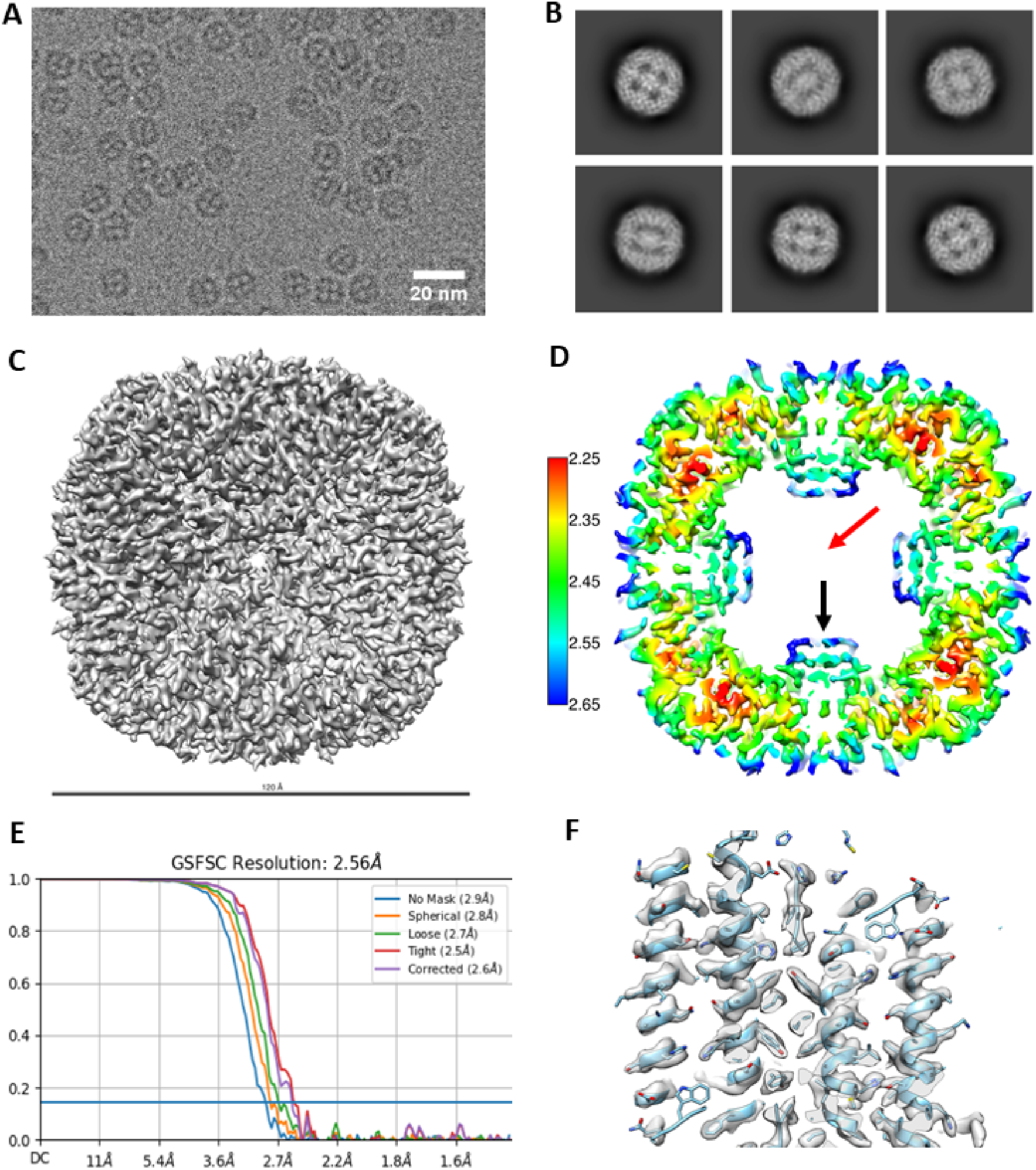
Data processing results for artemin-His. A) Representative micrograph of His tagged artemin showing the central cavity is distinctly filled with density attributed to 6xHis tags on each monomer. B) 2D classes showing that while the C-term of the monomer does point inwards from the shell, and fully fills the central cavity. C) Cryo-EM map of artemin-His with ∼120 Å diameter D) Thin virtual slice through a resolution heat map showing the C-term alpha helices pointing inwards (black arrow) into the cavity. Central cavity does not show any density for His tags corresponding to the 2D classes or micrographs (red arrow). Scalebar indicates resolution in Å. E) Resolution estimated by gold standard at 2.56 Å at 0.143 FSC. F) Quality of the map as inspected by fitting of alpha helices.

Previous biochemical studies combined with homology modeling have indicated that several conserved cysteines in artemin are essential for structural integrity and the putative chaperone activity of artemin ^8^ while the C-terminus was found to be important for the overall thermostability of artemin. Additionally, recent reports have identified the artemin dimer as the putative unit that has chaperone activity. In our experimentally derived model, the artemin dimer is oriented similarly to an apoferritin dimer **(Figure 2 B)** and a disulfide bridge exists between Cys61 and Cys22 of neighboring opposite facing monomers. This confirms the presence of 2 disulfide bridges per dimer **(Figure 2 C)** which is in line with previous homology modeling ^8^ and biochemical studies ^9^ that identified structural but not functional artemin destabilization at high temperatures when either or both of these Cys residues were modified. None of the other 8 cysteines are seen to be involved in disulfide bridges although all are surface exposed.

In addition, the overall octahedral symmetry shows an extra stabilizing interaction where the L’ loops or two monomers form a hashtag arrangement that connects the 4 helix bundles from each monomer in addition to the ferritin-like L loop interaction between two monomers at the outer surface **(Figure 2 D)**. Somewhat surprisingly, loop L’ and helix F extend and contact neighboring dimers which differs from all prior reported homology modeling efforts. This results in helices F from each monomer forming a second 4-helix arrangement toward the center of the complex **(Figure 2 D-F)**. Helix E from one dimer interacts with helices from 3 neighboring dimers around a 4-fold axis similar to apoferritin. For example, the ferritin monomer would contact chains at the 2 (dimer), 3 and 4-fold interface. In addition, near the 4-fold interface in artemin, helices F from neighboring chains form a second interaction facilitated by the respective antiparallel loops L’ **(Fig 2A, E)** These additional inter- and intradimer interactions resulting from Loop L’ and helix F may contribute to the significant thermal stability of artemin.

Models of artemin created with AlphaFold2 and RosettaFold (**Supplementary Figure 4**) ^26,27^ show a similar fold for the core region (as expected due to high homology with ferritin), but they fail to capture the C-terminal Loop L’ and full Helix-F positioning. The hashtag arrangement and the interactions of Helix F with the neighboring dimer may be important in the context of the 24mer structure, they may rearrange when exposed to temperature or oxidation when in their dimer or monomer state and these may be what AlphaFold2 and RosettaFold are predicting. Though further experimental work will need to be performed to validate those models.

**Figure 4.**
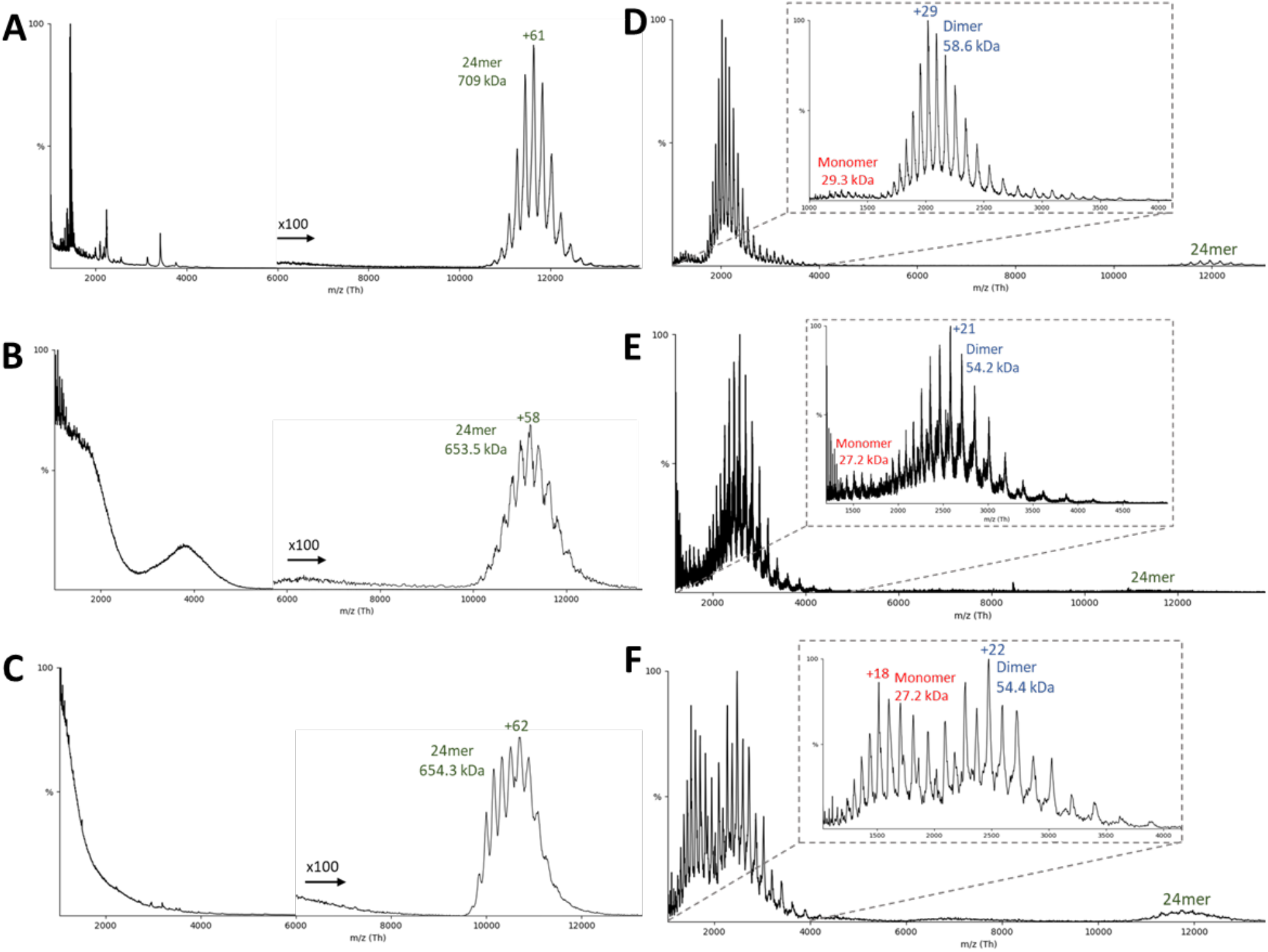
Native mass spectrometry of artemin constructs. (A-C) Representative native MS spectrum and the corresponding (D-F) collision induced dissociation (CID) spectrum of the resulting released monomers/dimers. (A/D) Flag-artemin, (B/E) artemin-His, and (C/F) Fluorescent artemin-His.

### 3.3 Structural perturbation identified potential features that affect artemin’s stability

The C-terminal helices of artemin are implicated in chaperone activities ^12^ and previous homology models suggested that the C-terminus fully packs the inside core of artemin. While our cryo-EM map of Flag-artemin clearly shows that the native C-terminus does not fully pack the interior of the artemin octahedral complex, we wondered what would happen if we intentionally filled that cavity with extra amino acids. We therefore purchased a second clone of artemin with a C-terminal 6xHis tag (artemin-His) that would permit possible filling of the inner cavity while also addressing whether one could purify artemin using a tag on the C-terminus. We were able to successfully express and purify artemin-His with similar yields as Flag-artemin. Based on biochemical analyses **(Supplementary Figure 1 C, D)**, we obtained a fully assembled 24mer of artemin-His despite the tag being putatively localized to the interior of the complex. Cryo-EM analyses and image processing revealed certain differences between the N- and C-terminal tagged constructs. First, the central cavity of the artemin-His appeared to be filled both in the micrographs and 2D class averages **(Figure 3 A, B)** as well as resulting 3D volume. This excess density relative to Flag-artemin is attributed to the 6xHis tag itself. A total of 192 amino acid residues (each monomer has a 2 amino acid linker and 6 His; 24 × 8) were added and these successfully filled the cavity **(Figure 3 C, D)**. However, no refined density was observed in the 3D map at the very center suggesting a lack of any discernable secondary structure in the 6x His tag. The final map obtained was at 2.56 Å (0.143 FSC) **(Figure 3 E, F)** was of lower resolution than the Flag-artemin map and a comparison of the C1 (no symmetry) versus octahedral symmetry map showed a minor disruption to the packing symmetry in the octahedral form which explains the lower resolution. We postulate that the minor disruption to the symmetry packing is due to at least one of the C-terminal 6xHis tags being excluded from the inner cavity due to full packing of all the other tags and exiting the complex through either the 3-fold or 4-fold channels. This is supported by the observation that affinity purification of intact octahedral complexes using the His tag at the supposedly buried C-terminus was attainable at similar yields as Flag-artemin purification and near 90% of total expressed artemin. While the C-terminal tag slightly affected the resolution and assembly symmetry of artemin, we wanted to also check the effect on stability using native MS which is not an image-based approach.

Using native MS, the mass of the intact 24mer for Flag-artemin was observed to be 709 kDa (theoretical: 696 kDa) while artemin-His was 653.5 kDa (theoretical: 650 kDa) **(Figure 4A-C)**. After isolating the 24mer, collision-induced dissociation (CID) was used to release smaller subunits **(Figure 4D-E)**. In CID, the protein ions are accelerated into a pressurized collision cell where the protein ions then collide with a neutral gas (argon in this experiment). As the number of collisions increase, the internal energy of the protein increases as well, causing potential unfolding and release of smaller subunits and/or bound ligands ^28,29^. Typically, a monomer is expected to be stripped from the complex during CID. In the case of the Flag-artemin **(Figure 4D)** and artemin-His **(Figure 4E)**, only a small population of monomers was observed, but the predominant species was dimers. This unusual CID behavior is consistent with the observed inter-subunit disulfide linkages in the cryo-EM structure (**Figure 2C**). In addition, we attempted to disrupt stability of the complex by doping fluorescent lysine tRNA into the cell-free reaction in hopes this approach might stress the complex assembly or dimer stability due to small steric hindrance. The use of doping rather than complete swapping of all lysine tRNA permitted the random incorporation of fluorescently labelled lysines into the artemin monomer. This was important since there are 15 lysines in the full-length 230 amino acid sequence of artemin (excluding tags) with one lysine being immediately adjacent to Cys61 involved in disulfide bonding and several in Loop L’ and at the C-terminus. The fluorescent artemin-His (Fluor artemin-His) complex expressed and purified like artemin-His and was found to be a clean octahedral complex by Native PAGE and was observed as a 24-mer by native MS at 654.3 kDa (**Figure 4C**). Interestingly, when CID was performed with the same settings as used above for Flag-artemin and artemin-His, nearly equivalent levels of monomeric and dimeric species were released from Fluor artemin-His **(Figure 4F)**. The masses of the released monomers and dimers in the Fluor artemin-His were essentially the same as those in the artemin-His, within experimental error, and cannot account for incorporation of any fluorescent tag. The presence of detectable monomeric species suggests that Fluor artemin-His disrupted part of the dimeric substructure, likely via prevention of inter-subunit disulfides. Additionally, Fluor artemin-His showed higher charge state distributions than the other artemin 24mers (**Figure 4C vs Figure 4A-B**). The charge state distribution for Fluor artemin-His also appeared to be less symmetric than the other two, suggesting multiple overlapping distributions (a more distinct bimodal distribution from another replicate is shown in **Supplementary Figure 5**). It is generally accepted that charge state distribution correlates with protein conformation, although the detailed mechanisms are under debate ^30,31^. The higher charge state of Fluor artemin-His imply a less compact structure potentially due to the disruption of the interfaces. Therefore, the change of charge state distributions for the different complexes indicates changes to the structures in response to terminal and lysine tagging.

## 4 Discussion

Here we describe a method that allowed us to progress from receiving a custom synthesized gene/plasmid, through expression, purification and cryo-EM structure determination at sub 2.5 Å resolution within 2 weeks. This is also first report of an experimental structure for the diapause chaperone artemin, almost 40 years after it was first discovered. We found that the C-terminal region important for chaperoning is positioned differently than all prior homology modeling, molecular dynamics and even recent Alphafold2 and RosettaFold models suggest. The C-terminal Loop L’ and Helix F were observed to provide additional interfaces for artemin dimers to interact and stabilize the 24mer assembly. These results raise new questions regarding the structural details of how artemin actually functions as a chaperone. For example, the functional chaperone unit of artemin is believed to be the dimer form but does it retain the same overall fold as the dimer in the 24mer or does the C-terminal region (or other regions) refold during chaperoning? A logical extension of our study would be structural studies of artemin “caught in the act” of chaperoning a target protein like citrate synthase or lysozyme. An artemin monomer or dimer on its own would be difficult to resolve using single particle cryo-EM, however a dimer interacting with the chaperoned target would be big enough for both native MS and cryo-EM studies, as long as the binding interfaces between artemin dimer and target are specific. Native MS methods such as collision induced unfolding ^32^ and variable temperature ^33^ electrospray will also provide unique contributions to the biophysical characterization of the stability and dynamics of these assemblies.

Molecular chaperones are broadly divided into holdases and foldases. Foldases are ATP dependent chaperones which actively support folding of proteins in the right conformation. Examples from bacteria include the GroEL/GroES or the DnaK/DnaJ/GrpE system; while the Hsp60/70/90 family of chaperones constitute the better studied foldases in mammalian systems ^34,35^. Holdases, also called small heat shock proteins (sHSPs) are ATP-independent chaperones. The bacterial protein Hsp33 is very well studied sHSP in bacteria, while the Get3 (yeast) and its human analog TRC40 are examples of holdases. Biochemical reports suggest that many holdases are regulated and reversibly activated via a redox switch. Brine shrimp have been reported to have their own complement of holdase (p26) and foldase (Hsc70) chaperones along with artemin ^1^. Foldases seem to prefer higher molecular weight assemblies (GroEL) while holdases typically exist as monomers or dimers of 10-40 kDa ^36^ and dimerize on stress dependent activation. In contrast to other holdases, artemin exists as a 24mer and upon exposure to stress, releases oligo n-mers of which dimers are most abundant. Artemin also lacks an α-crystallin domain which is otherwise ubiquitous in sHSPs but does form head to tail dimers like 2-Cys perioxiredoxins (2-Cys Prxs) – another redox mediated holdase ^37^. In contrast to artemin, upon exposure to increasing amount of stress, 2-Cys Prxs forms higher molecular weight assemblies (10 or 12 mers). The most drastic difference between artemin and other holdases is the irreversible structural changes that occur on exposure to stresses like heat or H_2_O_2_ whereas other redox regulated holdases are reversible ^13^. Artemin therefore appears to be a holdase-like chaperone with unique properties; especially since it acts as both a protein and RNA chaperone. With the structure of artemin now solved in the native 24mer state, it should be possible with future studies to dissect the structural basis and molecular mechanisms behind its RNA and protein chaperoning activity.

## 5 Contribution to the Field

While cryo-EM is routinely used to obtain high-resolution structures of macromolecules, the technique usually relies on homogenous, monodisperse and highly pure samples. Even the best-case scenarios typically involve several months for optimizing protein expression and purification, sample preparation and parameters for cryo-EM imaging. Here we present a workflow that allowed us to express, purify and solve a high-resolution cryo-EM protein structure in less than 2 weeks after receiving the gene clone. We coupled the use of a cell-free protein expression system to native MS and electron microscopy to solve the 2.04 Å structure of full-length artemin – a diapause chaperone protein with some homology to ferritin but which lacks a prior experimentally determined structure. We also investigated the stability of the complex 24-mer, dimer and monomer states and our atomic model clearly shows why artemin is unable to sequester iron as Loop L’ occludes access to critical residues even if they were mutated to consensus for ferritin. Future research on the mechanism of artemin chaperoning action for both RNA and protein protection, which hitherto relied on only *in silico* models based off homology to ferritin, will benefit from the high-resolution structure presented here.

## 7 Author Contributions

JEE conceived the research. JTP performed all cell-free expression and purification experiments. ADP performed all cryo-EM imaging and data analysis. SMP helped with protein modeling, performed native mass spectrometry experiments and data analysis with help from MZ. THM and INV assisted with cryo-EM data collection and model fitting respectively. ADP, SMP and JEE wrote the first draft of the manuscript and all authors edited the manuscript.

## 9 Acknowledgments

The research was performed using EMSL (grid.436923.9), a DOE Office of Science User Facility sponsored by the Biological and Environmental Research program located at PNNL.

## 11 Supplementary Material

Supplementary Material contains additional text and figures supporting this manuscript.

## 12 Data Availability Statement

The datasets for this study can be found at the PDB and EMDB repositories with ascension numbers EMD-24706, EMD-24707 and PDB:7RVB, to be released with publication.

