## Supplementary Material for "Cryo-EM structure of the diapause chaperone artemin"

### 1 Supplementary Figures and Tables

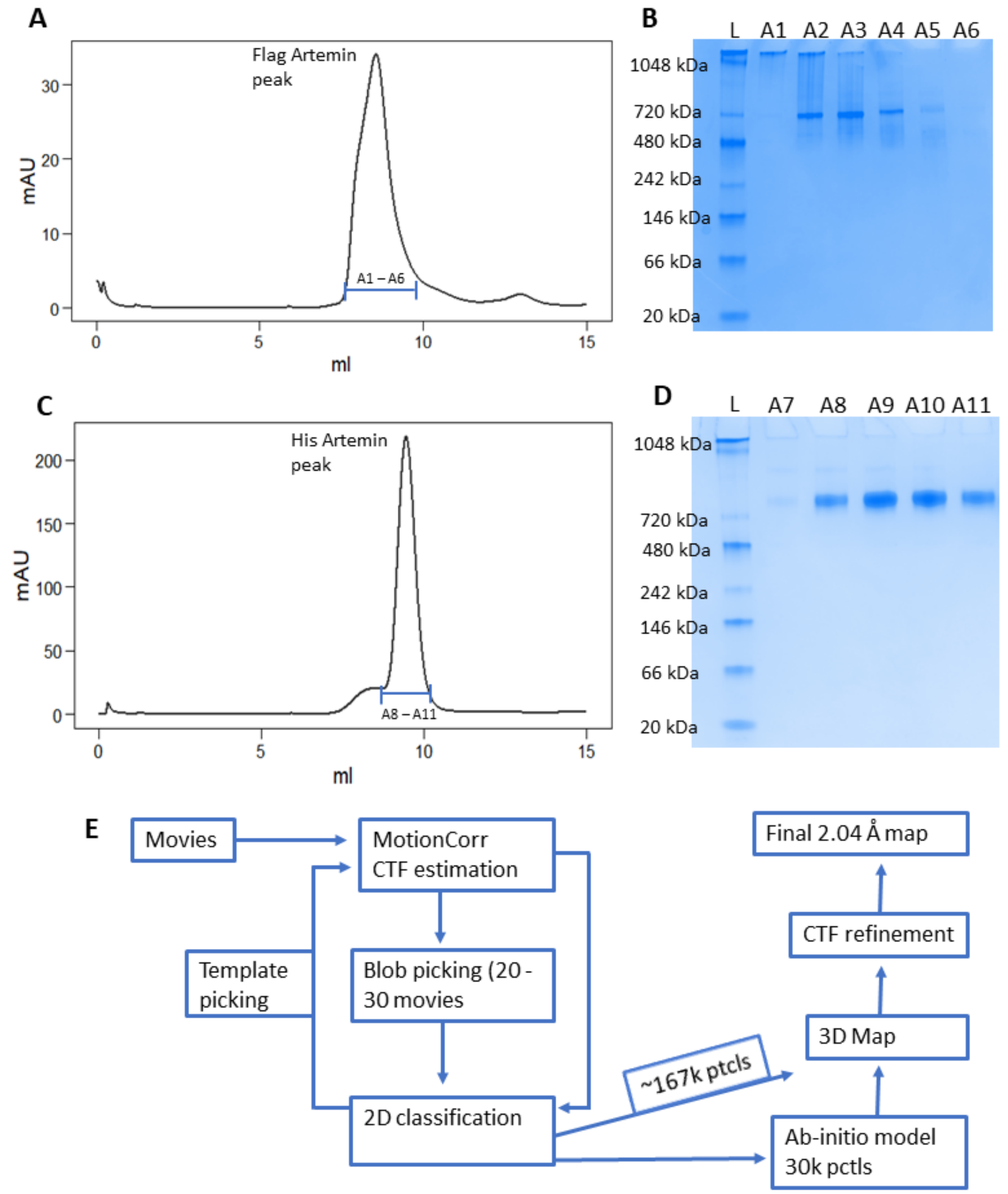


**Supplementary Figure 1 – Biochemistry and image processing workflow for Flag-artemin.** A) FPLC trace of artemin purified over a Superdex 200 column with a single homogenous peak for artemin. Gel B) Native PAGE analyses of fractions A1-A6 from the FPLC. Distinct clear bands near ~720 kDa mark indicate a 24mer of Flag-artemin. C) FPLC trace of artemin purified by SEC with a single homogenous peak for artemin. D) Native PAGE analyses of fractions A7-A11 from the FPLC. Distinct clear bands near ~720 kDa mark indicate a 24mer of Flag-artemin. L = lane for NativeMark Protein Ladder. E) Image processing workflow implemented using cryoSPARC and cryoSPARC Live.

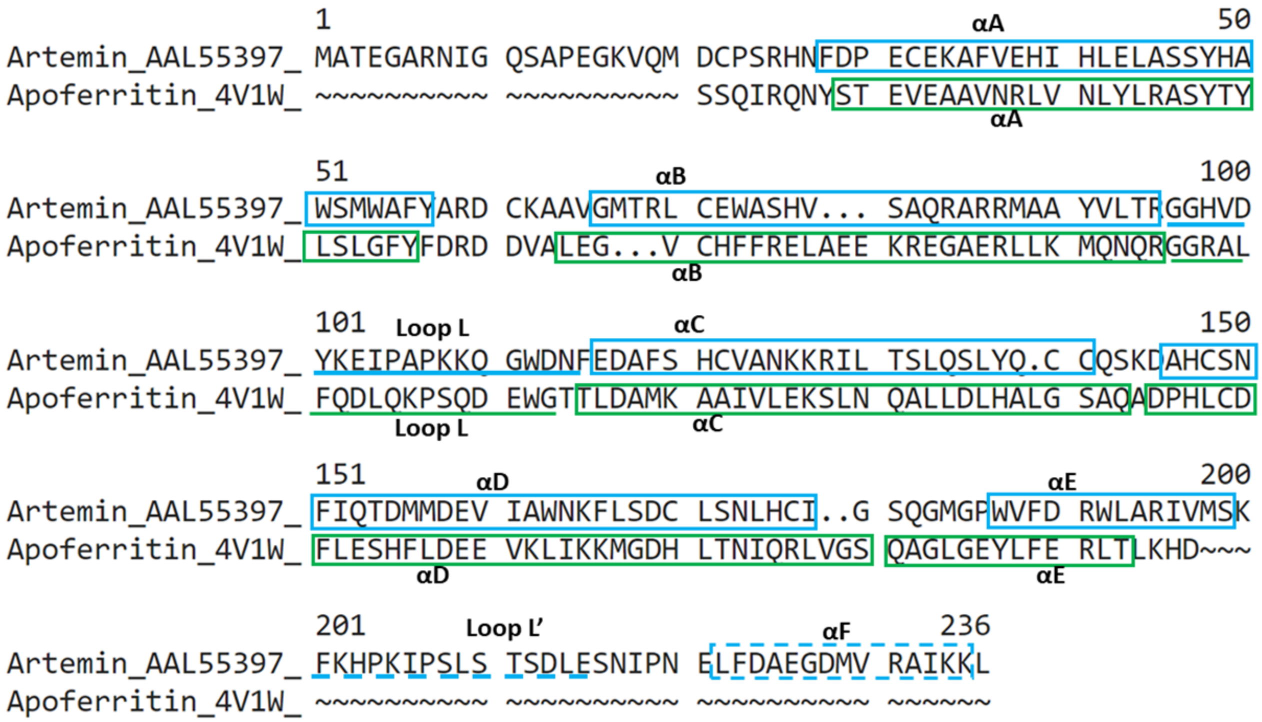


**Supplementary Figure 2 – Secondary structure based sequence comparison between artemin and apoferritin.** Primary sequences of artemin and apoferritin (4V1W) are aligned and the known alpha helical regions and loops are compared regions. Artemin has 6 α-helices (A-F, blue border) and 2 Loops (L, L' blue line) relative to 5 α-helices (A-E, green boundary) and 1 loop (green line). The Loop L' and Helix F in artemin indicated in dashed lines have no comparative region in apoferritin.

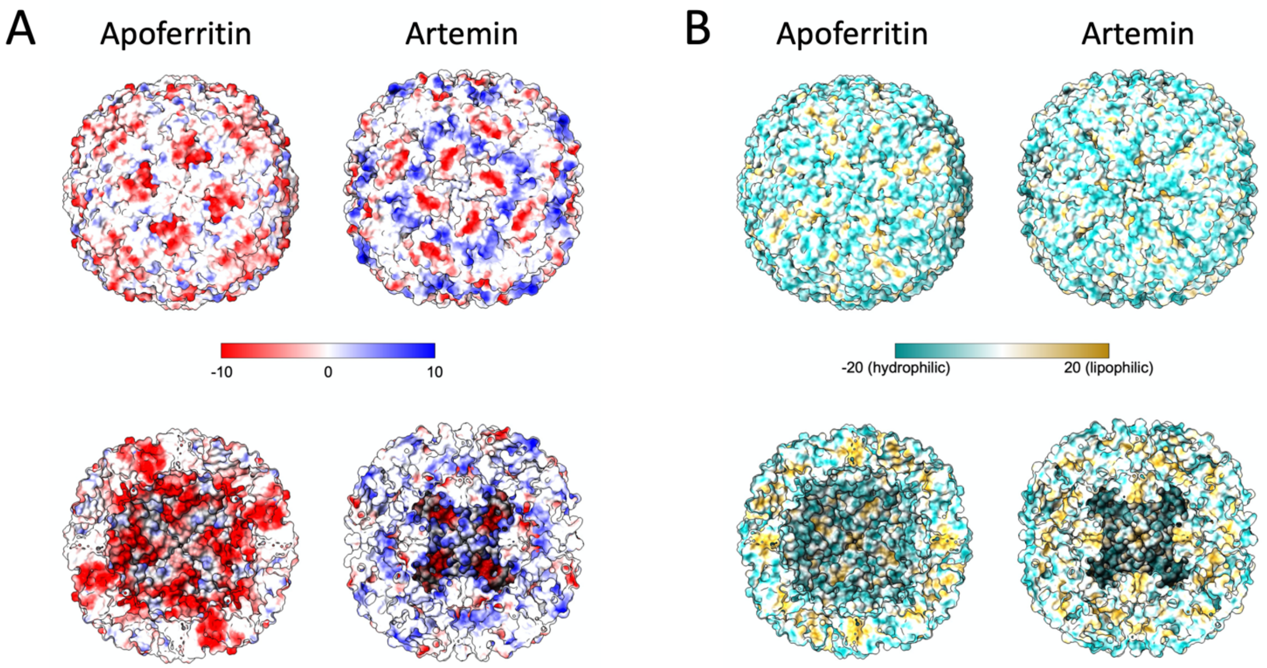


**Supplementary Figure 3 – Comparing electrostatic and coulombic surface maps of apoferritin and artemin.** A) Electrostatic charges visualized on the overall map of apoferritin and artemin (scale -10 to 10) as visualized in UCSF ChimeraX. Artemin has relatively more positive (blue) charge on surfaces exposed to the solvent externally and internally. B) Coulombic (hydrophobicity) map of apoferritin and artemin (scale -20 to 20) as visualized in UCSF ChimeraX. For both panels, the top images show the outer surface map while the bottom images show the internal view cutting halfway through the assembled 24mer. Artemin has more hydrophilic residues (cyan) exposed on the surface. All surface rendering views aligned along the C4 axis.

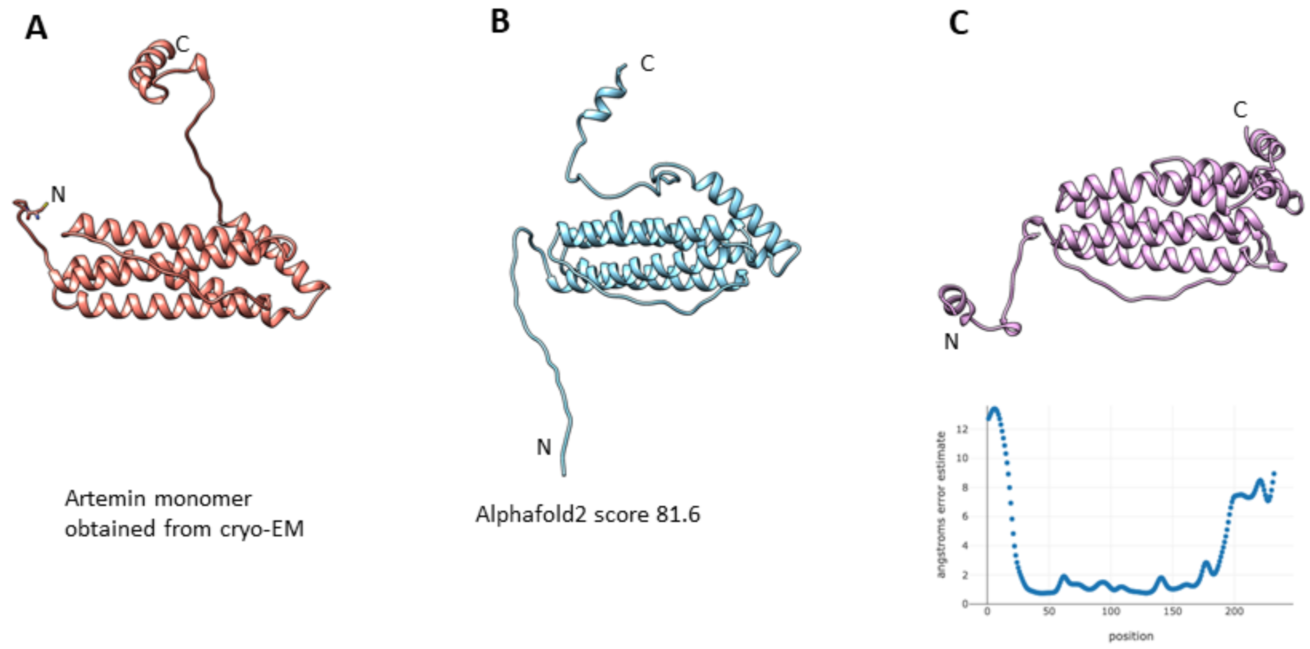


**Supplementary Figure 4 – Comparison of the structure of artemin monomer from with models obtained from Alphafold2 and RosettaFold.** A) A monomer of artemin calculated by direct refinement and fitting within our cryo-EM map, with its N to C orientation shown (black). B&C) Artemin monomer models calculated with AlphaFold2 and RosettaFold, respectively.

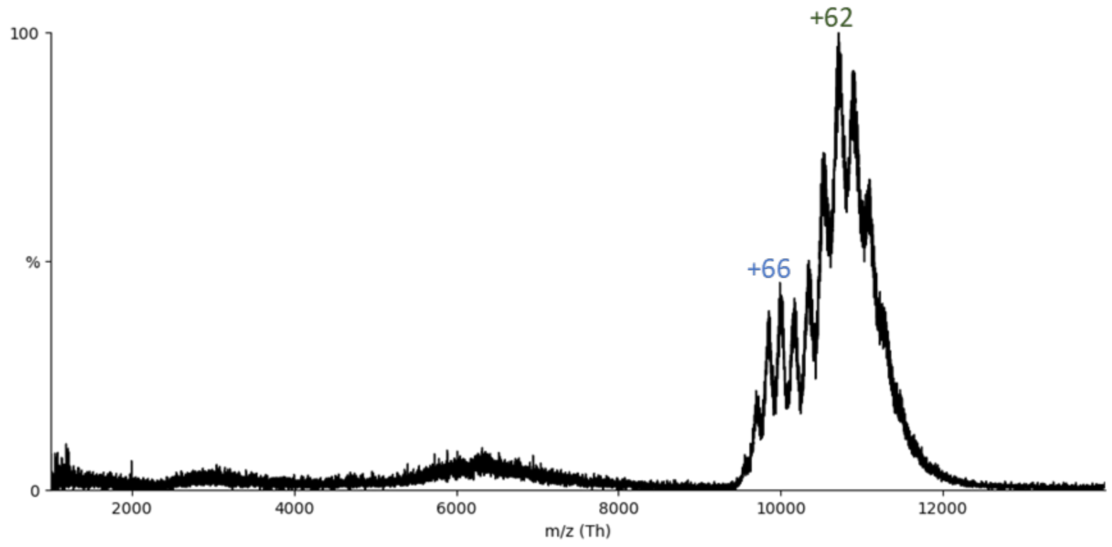


**Supplemental Figure 5 – Unfolding of fluorescent artemin-His.** Native MS shows two distinct charge state populations centered around +66 and +62, indicating the multiple conformations (potentially less compact at +66) of Fluor artemin-His.

**Supplement Table 1 – Amino acid sequence for all clones**

| | |
|---|---|
| **artemin-His** | MATEGARNIGQSAPEGKVQMDCPSRHNFDPECEKAFVEHIHLELASSY HAWSMWAFYARDCKAAVGMTRLCEWASHVSAQRARRMAAYVLTR GGHVDYKEIPAPKKQGWDNFEDAFSHCVANKKRILTSLQSLYQCCQS KDAHCSNFIQTDMMDEVIAWNKFLSDCLSNLHCIGSQGMGPWVFDR WLARIVMSKFKHPKIPSLSTSDLESNIPNELFDAEGDMVRAIKKL**GSHH HHHH** |
| **Flag-artemin** | **MDYKDHDGDYKDHDIDYKDDDDKL**ATEGARNIGQSAPEGKVQMDC PSRHNFDPECEKAFVEHIHLELASSYHAWSMWAFYARDCKAAVGMT RLCEWASHVSAQRARRMAAYVLTRGGHVDYKEIPAPKKQGWDNFED AFSHCVANKKRILTSLQSLYQCCQSKDAHCSNFIQTDMMDEVIAWNK FLSDCLSNLHCIGSQGMGPWVFDRWLARIVMSKFKHPKIPSLSTSDLES NIPNELFDAEGDMVRAIKKL |

- Sequences for affinity tags (6xHis and 3xFlag) along with spacers are shown bold and underlined

**Supplement Table 2 – Dataset details for all artemin constructs**

| **Dataset / EMDB ID** | **artemin-His EMD-24707** | **Flag-artemin EMD-24706** |
|---|---|---|
| Magnification | 130,000 | 130,000 |
| Super resolution | Yes | Yes |
| Pixel size (Å) | 0.3398 | 0.3398 |
| Total dose ($e^-$/ Å$^2$) | 52.33 | 41 to 58.59 |
| Defocus range (µm) | -0.3 to -1.3 | -0.3 to -1.3 |
| Micrographs collected | 6578 | 6626 |
| Micrographs utilized | 2494 | 4933 |
| Initial particles no. | 792,493 | 964,141 |
| Final particle no. | 660,454 | 167,408 |
| Symmetry imposed | Octahedral | Octahedral |
| Resolution (0.143 FSC) | 2.58 Å | 2.04 Å |
| Concentration loaded | 1 mg/ml | 1.5 mg/ml |

**Supplement Table 3 – Model validation statistics for artemin 24mer (PDB: 7RVB)**

| | |
|---|---|
| **Composition** | |
| Chains | 24 |
| Atoms | 40224 (Hydrogens: 0) |
| Residues | Protein: 5016, Nucleotide: 0 |
| Water | 0 |
| Ligands | 0 |
| **Bonds (RMSD)** | |
| Length (Å) (# > 4s) | 0.004 (0) |
| Angles (°) (# > 4s) | 0.543 (3) |
| Molprobity score | 1.12 |
| Clashscore | 1.94 |
| **Ramachandran plot (%)** | |
| Outliers | 0.00 |
| Allowed | 2.90 |
| Favored | 97.10 |
| Rotamer outliers (%) | 0.55 |
| Cß outliers (%) | 0.00 |
| **Model vs. Data** | |
| CC (volume) | 0.91 |

**Supplement Table 4 – Helices and loops of artemin based on similarity to apoferritin helices**

| **Residue number** | **Description** |
|---|---|
| 30-57 | Helix A |
| 65-92 | Helix B |
| 93-111 | Loop L |
| 112-137 | Helix C |
| 142-173 | Helix D |
| 179-192 | Helix E |
| 193-208 | Loop L' (not present in apoferritin) |
| 216-229 | Helix F (not present in apoferritin) |
